# Identification of the CoA-ester intermediates and genes involved in the cleavage and degradation of the steroidal C-ring by *Comamonas testosteroni* TA441

**DOI:** 10.1101/2021.06.08.447645

**Authors:** Masae Horinouchi, Toshiaki Hayashi

## Abstract

*Comamonas testosteroni* TA441 degrades steroids aerobically via aromatization of the A-ring accompanied by B-ring cleavage, followed by D- and C-ring cleavage. We previously revealed major enzymes and intermediate compounds in A,B-ring cleavage, β-oxidation cycle of the cleaved B-ring, and partial C,D-ring cleavage process. Here, we elucidated the C-ring cleavage and the β-oxidation cycle that follows. ScdL1L2, a 3-ketoacid Coenzyme A (CoA) transferase which belongs to the SugarP_isomerase superfamily, was thought to cleave the C-ring of 9-oxo-1,2,3,4,5,6,10,19-octanor-13,17-secoandrost-8(14)-ene-7,17-dioic acid-CoA ester, the key intermediate compound in the degradation of 9,17-dioxo-1,2,3,4,10,19-hexanorandrostan-5-oic acid (3aα-*H*-4α [3′-propionic acid]-7aβ-methylhexahydro-1,5-indanedione; HIP)-CoA ester in the previous study; however, this study suggested that ScdL1L2 is the isomerase of the derivative with a hydroxyl group at C-14 which cleaves C ring. The subsequent ring-cleaved product was indicated to be converted to 4-methyl-5-oxo-octane-1,8-dioic acid-CoA ester mainly by ORF33-encoded CoA-transferase (named ScdJ), followed by dehydrogenation by ORF21 and 22-encoded acyl-CoA dehydrogenase (named ScdM1M2). Then a water molecule is added by ScdN for further degradation by β-oxidation. ScdN is considered to catalyze the last reaction in C,D-ring degradation by the enzymes encoded in the steroid degradation gene cluster *tesB* to *tesR*.

**IMPORTANCE:** Studies on bacterial steroid degradation were initiated more than 50 years ago primarily to obtain materials for steroid drugs. Steroid-degrading bacteria are globally distributed, and the role of bacterial steroid degradation in the environment as well as in human is attracting attention. The overall degradation of steroidal four rings is proposed, however there are still much to be revealed to understand the complete degradation pathway. This study aims to uncover the whole steroid degradation process in *C. testosteroni*, which is one of the most studied representative steroid degrading bacteria and is suitable for exploring the degradation pathway because the involvement of degradation-related genes can be determined by gene disruption.

## INTRODUCTION

Actinobacterium *Rhodococcus equi* (formerly *Nocardia restrictus*) and proteobacterium *Comamonas testosteroni* (formerly *Pseudomonas testosteroni*) are known for their ability to degrade steroid compounds and their testosterone degrading mechanism was extensively studied for the primary purpose of obtaining materials to synthesize steroidal drugs around 1960. The research led to the identification of the major intermediate compounds in the A- and B-ring degradation processes, indicating that the steroidal degradation in actinobacterium and proteobacterium were similar [1–10]. Genetic studies on degradation of bacterial steroids started around the year 1990 in *C. testosteroni*, and the enzymes catalyzing the early steps of steroidal degradation (17β-dehydrogenase, 3α-dehydrogenase, 3-oxo-Δ5-steroid isomerase, Δ1-dehydrogenase, and Δ4-dehydrogenase) were identified [11–27]. However, the enzymes for ring-cleavage process remained unclear when we started investigating steroidal degradation in *C. testosteroni* TA441 in 1999 to reveal the precise mechanism of steroidal degradation as the leading model of bacterial aerobic steroid degradation. The mechanism of steroidal degradation in TA441 is shown in Fig. 1. TA441 degrades steroids (e.g. testosterone, cholic acid, and their derivatives) to androsta-1,4-diene-3,17-dione (ADD) (R1,R2=H) or the corresponding derivative (cf. 7α,9α-dihydroxy-androsta-1,4-diene-3,17-dione in cholic acid degradation) (compound **I**, Fig. 1), then into 2-hydroxyhexa-2,4-dienoic acid (**VI**) and 9,17-dioxo-1,2,3,4,10,19-hexanorandrostan-5-oic acid (**VII**) (3aα-*H*-4α [3′-propionic acid]-7aβ-methylhexahydro-1,5-indanedione; HIP) or the corresponding derivative (cf. 7α,9α-dihydroxy-17-oxo-1,2,3,4,10,19-hexanorandrostan-5-oic acid in cholic acid degradation) via aromatization of the A-ring and the subsequent ring cleavage and hydrolysis [28–35]. Then, Coenzyme A (CoA) is incorporated into **VII** by ScdA [36], and the resulting **VII**-CoA ester is further degraded, mainly by two cycles of β-oxidation on the cleaved B-ring and on the cleaved C,D-rings [37–42]. The first cycle of β-oxidation involves the removal of two carbons of cleaved B-ring from **VII**-CoA ester to generate 9α-hydroxy-17-oxo-1,2,3,4,5,6,10,19-octanorandrostan-7-oic acid (**XII**) -CoA ester. Then the C-ring is dehydrogenated to produce 9,17-dioxo-1,2,3,4,5,6,10,19-octanorandrost-8(14)-en-7-oic acid (**XIV**) -CoA ester, which is followed by D-ring cleavage. The derivatives of intermediate compounds after D-ring-cleavage, 9-oxo-1,2,3,4,5,6,10,19-octanor-13,17-secoandrost-8(14)-ene-7,17-dioic acid (**XV**), 9-oxo-1,2,3,4,5,6,10,19-octanor-13,17-secoandrost-8(14)-en-17-oic acid (**XVa**), 9-oxo-1,2,3,4,5,6,10,19-octanor-13,17-secoandrosta-8(14),15-dien-17-oic acid (**XVb**), 13-hydroxy-9-oxo-1,2,3,4,5,6,10,19-octanor-13,17-secoandrost-8(14)-en-17-oic acid (**XVIa**), and 9-hydroxy-1,2,3,4,5,6,10,19-octanor-13,17-secoandrosta-13,15-dien-17-oic acid (**XVIIb**) were isolated from the culture of ScdL1^-^L2^-^ mutant and were identified with NMR, high-resolution MS (HRMS), and other analysis techniques when necessary (Fig. 1). Based on these compounds, **XV**-CoA ester, **XVI**-CoA ester, and **XVII**-CoA ester were proposed as intermediate compounds with only the C-ring remaining unaffected in steroidal degradation because it was isolated much more in amount than other two compounds [41]; yet the substrate of the ring cleavage by ScdL1L2 is not entirely clear. 4-Methyl-5-oxo-octane-1,8-dioic acid (**XX**) and 4-methyl-5-oxo-oct-3-ene-1,8-dioic acid (**XXa**) are identified as compounds produced after C-ring cleavage [42], suggesting that 6-methyl-3,7-dioxo-decane-1,10-dioic acid (**XIX**)-CoA ester is the C-ring cleavage product. **XIX**-CoA ester is presumed to be produced by C-ring cleavage of 4-hydroxy-9-oxo-1,2,3,4,5,6,10,19-octanor-13,17-secoandrostane-7,17-dioic acid (**XVIII**) -CoA ester, but **XVIII** has not been isolated in the previous studies. Therefore, C,D-ring cleavage process requires further investigation. Most of the steroidal degradation genes in TA441 are encoded in two clusters, located on both ends of the 120 kb mega-cluster of steroidal degradation genes (Fig. 1).

**Fig 1.**
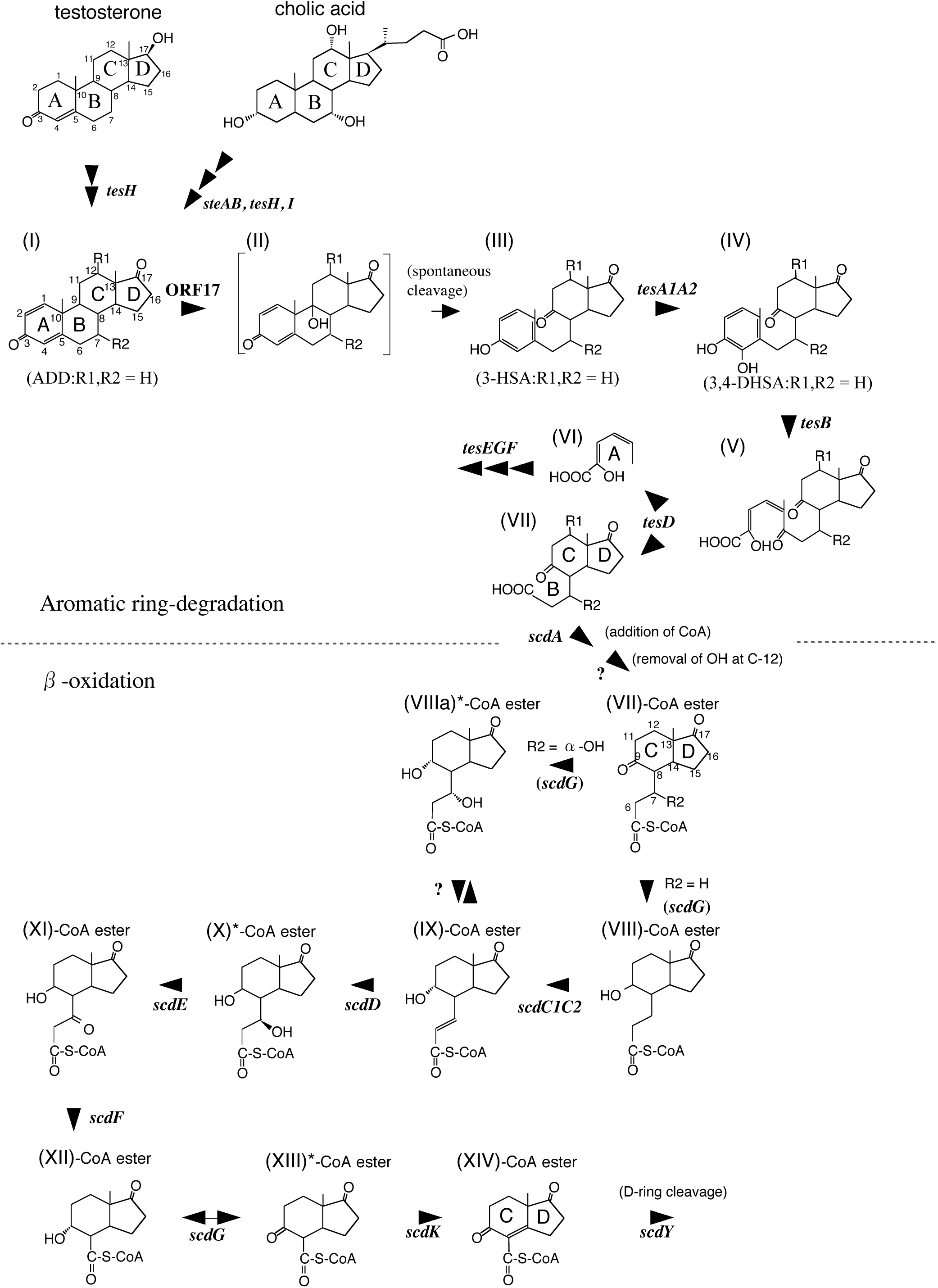
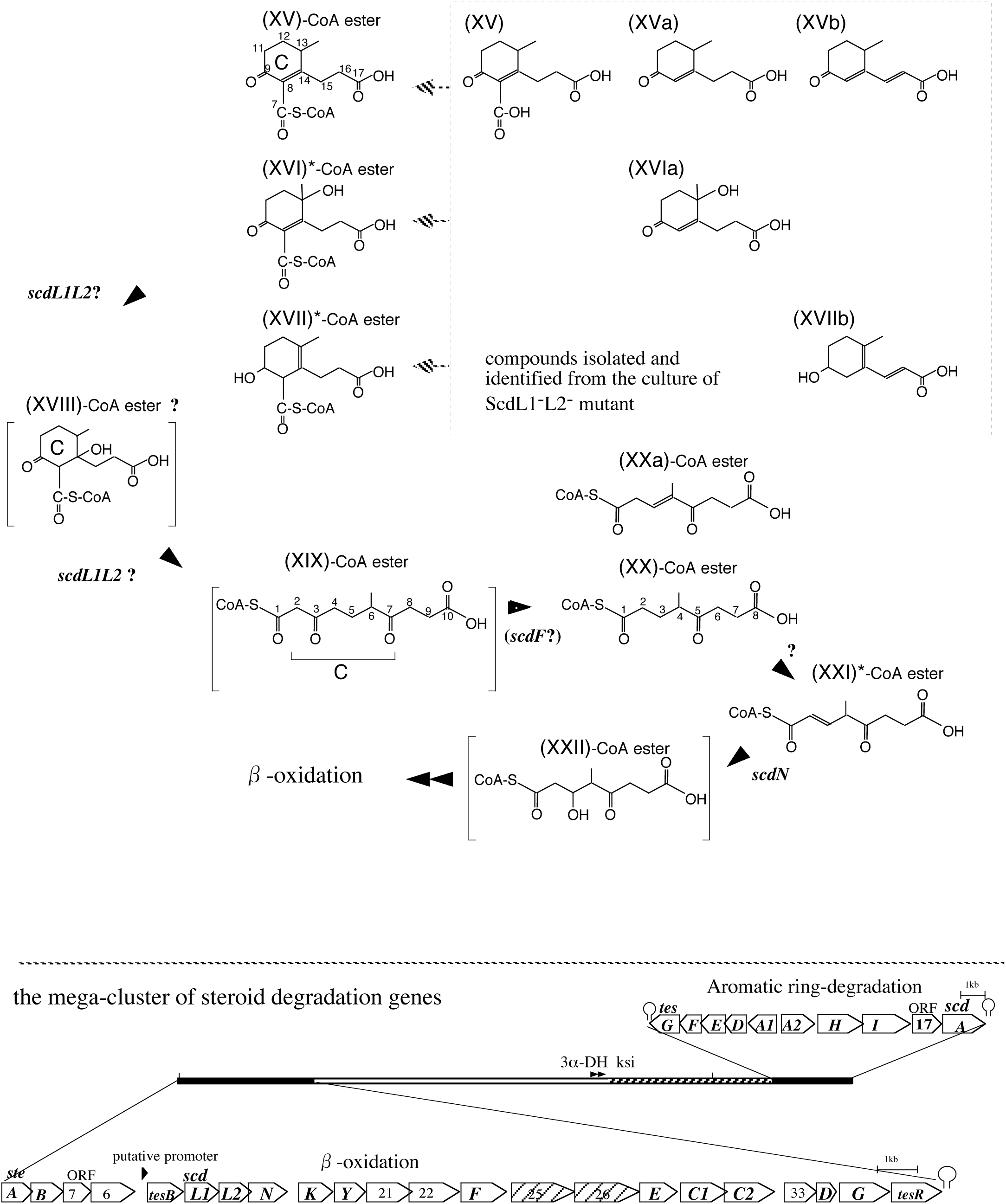
Steroid degradation pathway of *Comamonas testosteroni* TA441 revealed in our previous studies. Intermediate compounds were isolated and identified with NMR and high-resolution MS (HRMS) analysis (in β-oxidation process, compounds without CoA were identified) except for; compounds with * was detected by LC/MS and experimentally confirmed, and compounds in square brackets are presumption. C-ring cleavage process is not clear at the moment, therefore all the possible intermediate compounds with only C-ring are presented. The mega-cluster of steroid degradation genes in *C. testosteroni* TA441 is shown below the degradation pathway; the aromatic ring-degradation gene cluster (*tesG* to *scdA*) and the β-oxidation gene cluster (*steA* to *tesR*) locate both ends of this 120kb-mega cluster. 3α-Hydroxy-dehydrogenase (3α-DH) gene and 3-ketosteroid Δ4-5 isomerase (ksi) gene are in the DNA region between the two clusters. Possible degradation genes for the side chain of cholic acid at C17 are found in the striped region. Compounds are; androsta-1,4-diene-3,17-dione (ADD) (R1,R2=H), (**I**); 9-hydroxy-androsta-1,4-diene-3,17-dione (ADD) (R1,R2=H), (**II**); 3-hydroxy-9,10-secoandrosta-1,3,5(10)-triene-9,17-dione (3-HSA) (R1,R2=H), (**III**); 3,4-dihydroxy-9,10-secoandrosta-1,3,5(10)-triene-9,17-dione (3,4-DHSA) (R1,R2=H), (**IV**); 4,5-9,10-diseco-3-hydroxy-5,9,17-trioxoandrosta-1(10),2-dien-4-oic acid (R1,R2=H), (**V**); (2*Z*,4*Z*)-2-hydroxyhexa-2,4-dienoic acid, (**VI**); 9,17-dioxo-1,2,3,4,10,19-hexanorandrostan-5-oic acid (3aα-*H*-4α [3′-propionic acid]-7aβ-methylhexahydro-1,5-indanedione, HIP) (R1,R2=H), (**VII**); 9α-hydroxy-17-oxo-1,2,3,4,10,19-hexanorandrost-6-en-5-oic acid, (**VIII**), 9α,7α-dihydroxy-17-oxo-1,2,3,4,10,19-hexanorandrost-6-en-5-oic acid, (**VIIIa**), 9-hydroxy-17-oxo-1,2,3,4,10,19-hexanorandrost-6-en-5-oic acid (**IX**), 9α,7β-dihydroxy-17-oxo-1,2,3,4,10,19-hexanorandrost-6-en-5-oic acid, (**X**), 9α-hydroxy-7,17-dioxo-1,2,3,4,10,19-hexanorandrost-6-en-5-oic acid, (**XI**), 9α-hydroxy-17-oxo-1,2,3,4,5,6,10,19-octanorandrostan-7-oic acid (**XII**), 9,17-dioxo-1,2,3,4,5,6,10,19-octanorandrostan-7-oic acid (**XIII**), 9,17-dioxo-1,2,3,4,5,6,10,19-octanorandrost-8(14)-en-7-oic acid (**XIV**), 9-oxo-1,2,3,4,5,6,10,19-octanor-13,17-secoandrost-8(14)-ene-7,17-dioic acid (**XV**), 13-hydroxy-9-oxo-1,2,3,4,5,6,10,19-octanor-13,17-secoandrost-8(14)-ene-7,17-dioic acid (**XVI**), 9-hydroxy-1,2,3,4,5,6,10,19-octanor-13,17-secoandrost-13-ene-7,17-dioic acid (**XVII**), 9-oxo-1,2,3,4,5,6,10,19-octanor-13,17-secoandrost-8(14)-en-17-oic acid (**XVa**), 9-oxo-1,2,3,4,5,6,10,19-octanor-13,17-secoandrosta-8(14),15-dien-17-oic acid (**XVb**), 13-hydroxy-9-oxo-1,2,3,4,5,6,10,19-octanor-13,17-secoandrost-8(14)-en-17-oic acid (**XVIa**), 9-hydroxy-1,2,3,4,5,6,10,19-octanor-13,17-secoandrosta-13,15-dien-17-oic acid (**XVIIb**), 14-hydroxy-9-oxo-1,2,3,4,5,6,10,19-octanor-13,17-secoandrostane-7,17-dioic acid (**XVIII**), 6-methyl-3,7-dioxo-decane-1,10-dioic acid (**XIX**), 4-methyl-5-oxo-octane-1,8-dioic acid (**XX**), 4-methyl-5-oxo-oct-3-ene-1,8-dioic acid (**XXa**), 3-hydroxy-4-methyl-5-oxo-oct-2-ene-1,8-dioic acid (**XXI**), and hydroxy-4-methyl-5-oxo-octane-1,8-dioic acid (**XXII**). Enzymes are; SteA (dehydrogenase for 12-αOH to 12-ketone) [57], SteB (hydrogenase for 12-ketone to 12β-OH) [57], TesH (Δ1-dehydrogenase), ORF17-encoded enzyme (**I-**hydroxylase at C9), TesA1A2 (**III-**hydroxylase at C4), TesB (*meta*-cleavage enzyme for **IV**), TesD (**V-**hydrolase), TesE (**VI-**hydratase), TesF (aldolase), TesG (acetoaldehyde dehydrogenase), ScdA (CoA-transferase for **VII**), ScdG (hydrogenase primarily for 9-OH of **XII-**CoA ester), ScdC1C2 (Δ6-dehydrogenase for **VIII-**CoA ester), ScdD (**IX-**CoA ester hydratase), ScdE (**X-**CoA ester dehydrogenase at C7), ScdF (**XI-**CoA ester thiolase), ScdK (Δ8-14-dehydrogenase for **XIII-**CoA ester), ScdY (likely to be **XIV-**CoA ester hydratase, but further investigation required), ScdL1L2 (CoA-transferase/isomerase involved in C-ring cleavage), and ScdN (**XXI-**CoA ester hydratase).

Recently, degradation of bacterial steroids was also reported in other bacterial genera, specifically *Mycobacterium* [43, 44], *M. tuberculosis* H37Rv [45], *Rhodococcus* [46], *Pseudomonas* [47], *Novosphingobium tardaugens* [48] (aerobic degradation), with anaerobic androgen degradation in *Sterolibacterium* [49], *Denitratisoma* sp. [50], and aerobic estrogen degradation in *Sphingomonas* [51]. C,D-ring degradation has been proposed in *M. tuberculosis* H37Rv [45], which is similar but not identical to that in TA441. Degradation of bacterial steroids is considered to be important especially in *M. tuberculosis* H37Rv because the *mce4* operon in *M. tuberculosis*, which encodes a cholesterol import system, is essential for persistence in the lungs of chronically infected animals and for the growth within interferon-gamma-activated macrophages [52]. Cholesterol catabolism and broader utilization of *M. tuberculosis* are also important for the pathogen maintenance in the host [43].

In this study, we describe the degradation of the C-ring cleaved metabolites through the second β−oxidation cycle for elucidating the process of aerobic degradation of bacterial steroids.

## RESULTS AND DISCUSSION

### Identification of essential genes involved in the C-ring cleavage and degradation

Among the genes in the steroid-degradation gene cluster *tesB* to *tesR* (Fig. 1), the function of ORFs 21, 22, and 33 in the steroidal degradation process was yet to be confirmed. Both ORF21- and ORF22- encoded proteins showed homology to acyl-CoA dehydrobenase (ACAD) superfamily proteins (BLAST search). ORF21-encoded protein was similar to acyl-CoA dehydrobenase CaiA, putative acyl-CoA dehydrobenases FadE6_17_26, and PimC, the large subunit of pimenoyl-CoA dehydrogenase. ORF22-encoded protein was similar to acyl-CoA dehydrobenase CaiA and PimD, the small subunit of pimenoyl-CoA dehydrogenase. ORF33-encoded enzyme showed the highest homology to thiolase/acetyl-CoA acetyltransferases (cd00751), whose actual substrate is unknown. Then, mutants in which ORF21, 22, and 33 were disrupted individually (ORF21^-^, ORF22^-^, and ORF33^-^ mutants, respectively) were constructed, incubated with cholic acid, and the culture was analyzed with ultra-performance liquid chromatography/mass spectrometry (UPLC/MS). ORF21^-^ and ORF22^-^ mutants accumulated a considerable amount of 4-methyl-5-oxo-octane-1,8-dioic acid (**XX**), suggesting that these genes are involved in degrading **XX**-CoA ester (Fig. 2A and B). **XX,** in which all the steroidal rings are cleaved, is one of the major metabolites that accumulate during steroidal degradation by TA441 [41]. **XX**-CoA ester is dehydrogenated to 3-hydroxy-4-methyl-5-oxo-oct-2-ene-1,8-dioic acid (**XXI**)-CoA ester, which is followed by hydration by ScdN (Fig. 2) [41]. The detected amount of **XX** in ORF33^-^ mutant culture was smaller than that in ORF21^-^ and ORF22^-^ mutants, and almost the same level as that of ScdN^-^ mutant (Fig. 2C and D). **XXI** was detected in the ORF33^-^ and ScdN^-^ mutants; however, it remained undetected in the ORF21^-^ and ORF22^-^ mutants.

**Fig 2.**
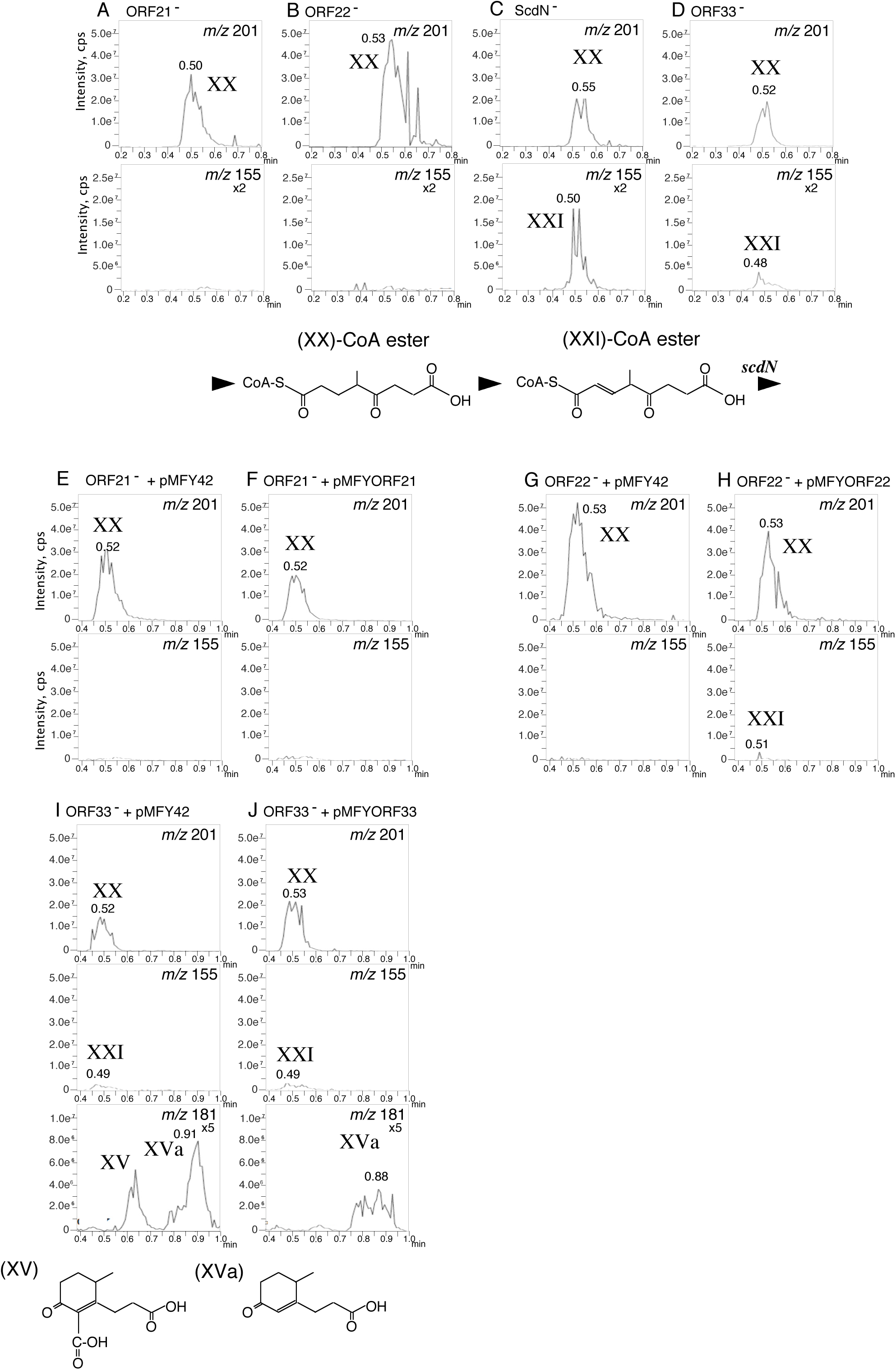
UPLC/MS analysis of the culture of the ORF21^-^, ORF22^-^, and ORF33^-^ mutants. Panels *m/z* 201 and 155 indicate the mass chromatogram of**XX** and **XXI**, respectively, in the culture of ORF21^-^ mutant (A), ORF22^-^ mutant (B), *scdN*^-^ mutant (C, as an authentic for **XX** and **XXI**), ORF33^-^ mutant (D), ORF21^-^ mutant carrying pMFY42 (E), ORF21^-^ mutant carrying pMFYORF21 (pMFY42-derivative carrying ORF21) (F), ORF22^-^ mutant carrying pMFY42 (G), ORF22^-^ mutant carrying pMFYORF22 (pMFY42-derivative carrying ORF22) (H), ORF33^-^ mutant carrying pMFY42 (I), and ORF33^-^ mutant carrying pMFYORF33 (pMFY42-derivative carrying ORF33) (J). Panels *m/z* 181 in Fig. 2I and 2J indicate **XV** (RT = 0.65 m) and **XVa** (RT = approximately 0.9 min). In mass chromatogram, the vertical axis indicates intensity (count/sec) and the horizontal axis indicates RT (min).

Then, we constructed an ORF21^-^mutant carrying the broad host range plasmid pMFY42 (2E, negative control), ORF21^-^ mutant carrying a pMFY42-based plasmid encoding ORF21 (pMFYORF21) (2F), ORF22^-^ mutant carrying pMFY42 (2G, negative control), ORF22^-^ mutant carrying a pMFY42-based plasmid encoding ORF22 (pMFYORF22) (2H), ORF33^-^ mutant carrying pMFY42 (2I, negative control), and ORF33^-^ mutant carrying a pMFY42-based plasmid encoding ORF33 (pMFYORF33) (2J). These mutants were incubated with cholic acid and the cultures were analyzed in the same manner. In the culture of ORF21^-^ mutant carrying pMFYORF21 and ORF22^-^ mutant carrying pMFYORF22, **XX** was smaller in amount than in ORF21^-^ and ORF22^-^ mutants carrying pMFY42, indicating ORF21 and ORF22 are involved in conversion of **XX**. The reduction in amount was smaller than we expected, probably because the expression level of each protein from pMFY42, a low copy plasmid, was not enough (cf. ScdY^-^ mutant carrying the pMFY42 with a copy of *scdY* showed a small reduction in quantity of **XIV** while the reduction in amount increased dramatically when we used ScdY^-^ mutant carrying the pMFY42 with multiple copies of *scdY* [39]). In contrast to ORF21^-^ and ORF22^-^ mutants, the amount of **XX** in the culture of ORF33^-^ mutant carrying pMFYORF33 was higher than that in ORF33^-^ mutant carrying pMFY42. ORF33^-^ mutant is the only mutant except for ScdL1^-^L2^-^ mutant that accumulates a considerable amount of **XVa** (9-oxo-1,2,3,4,5,6,10,19-octanor-13,17-secoandrost-8(14)-en-17-oic acid) among the gene-disrupted mutants of *tesB* to *tesR,* respectively (Fig. 2I *m/z* 181, Fig. S1D *m/z* 181). The amount of **XVa** in the culture of ORF33^-^ mutant carrying pMFYORF33 was smaller than that in the culture of ORF33^-^ mutant carrying pMFY42 (Figs. 2I and J, *m/z* 181). Since the homology search strongly suggested that the ORF33 encoded-enzyme is a thiolase/acetyl-CoA acetyltransferase, in combination with the aforementioned results, the ORF33 encoded-enzyme was considered to be involved in conversion of 6-methyl-3,7-dioxo-decane-1,10-dioic acid (**XIX**)-CoA ester to **XX**-CoA ester (Fig. 1). **XX** was detected in the culture of the ORF33^-^ mutant, possibly because ScdF also acts on **XIX**-ester (cf. Fig. 1 and the section “CoA-transferase, ScdF”). We named ORF33-encoded enzyme ScdJ; however, further analysis is required to confirm its function.

### Structural elucidation of the CoA-esters involved in the degradation of steroidal C-ring (I); analysis of compounds XX, XXa (4-methyl-5-oxo-oct-3-ene-1,8-dioic acid), and XXI using reverse-phase liquid chromatography with tandem mass spectrometry (LC/MS/MS)

**XX** and **XXa** were isolated from the culture of ScdN^-^ mutant, and identified via both high-resolution MS (HRMS) and NMR analysis in the previous study [41]. However, **XXI** was decarboxylated during the isolation procedure and UPLC/MS analysis only detected the decarboxylated form of **XXI** as a peak with an *m/z* of 155. The compound structure was confirmed with several experiments, however we did not successfully detect **XXI**, a peak with an *m/z* of 199 via UPLC/MS. Therefore, we performed LC/MS/MS analysis to directly confirm the presence of **XXI** in the culture.

**XX** and **XXa** were detected in the culture of the ORF21^-^22^-^ mutant, in which both ORF21 and 22 were disrupted, incubated with cholic acid. **XX** was detected as a peak with an *m/z* of 201 at retention time (RT) = 5.50 min with major fragments with *m/z* values of 183 (M^-^ - H_2_O), 157 (M^-^ - CO_2_^·^), 139 (M^-^ -CO_2_^·^ and - H_2_O), 129 (M^-^ - CH_2_CH_2_CO_2_^·^), 111 (M^-^ - CH_2_CH_2_CO_2_^·^ and H_2_O), and 85 (M^-^ - CH_2_CH_2_CO_2_^·^ and - CO_2_^·^) (Fig. 3A, mass chromatogram *m/z* 201). **XXa** was detected as a peak with an *m/z* of 199 at RT = 6.30 min with major fragments with *m/z* values of 181 (M^-^ - H_2_O), 155 (M^-^ - CO_2_^·^), 137 (M^-^ -CO_2_^·^ and - H_2_O), 127 (M^-^ - CH_2_CH_2_CO_2_^·^), 111 (M^-^ −2 CO_2_^·^), 109 (M^-^ - CH_2_CH_2_CO_2_^·^ and H_2_O), and 83 (M^-^ - CH_2_CH_2_CO_2_^·^ and - CO_2_^·^) (Fig. 3B, mass chromatogram *m/z* 199 at RT = 6.30 min). Furthermore, **XXI** was detected by LC/MS/MS as a peak with an *m/z* of 199 at RT = 5.45 min with major fragments with *m/z* values of 155 and 111 (Fig. 3C, mass chromatogram *m/z* 199 and *m/z* 155 at RT = 5.45 min, a peak of *m/z* 199 is not visible in the mass chromatogram). The mass spectrum data confirmed the structure of **XXI**, which had only been experimentally confirmed in the previous study. In addition, all the three compounds were detected more clearly on the chromatogram obtained using LC/MS/MS compared to the one obtained from UPLC/MS. Therefore, LC/MS/MS was used in the subsequent analyses.

**Fig 3.**
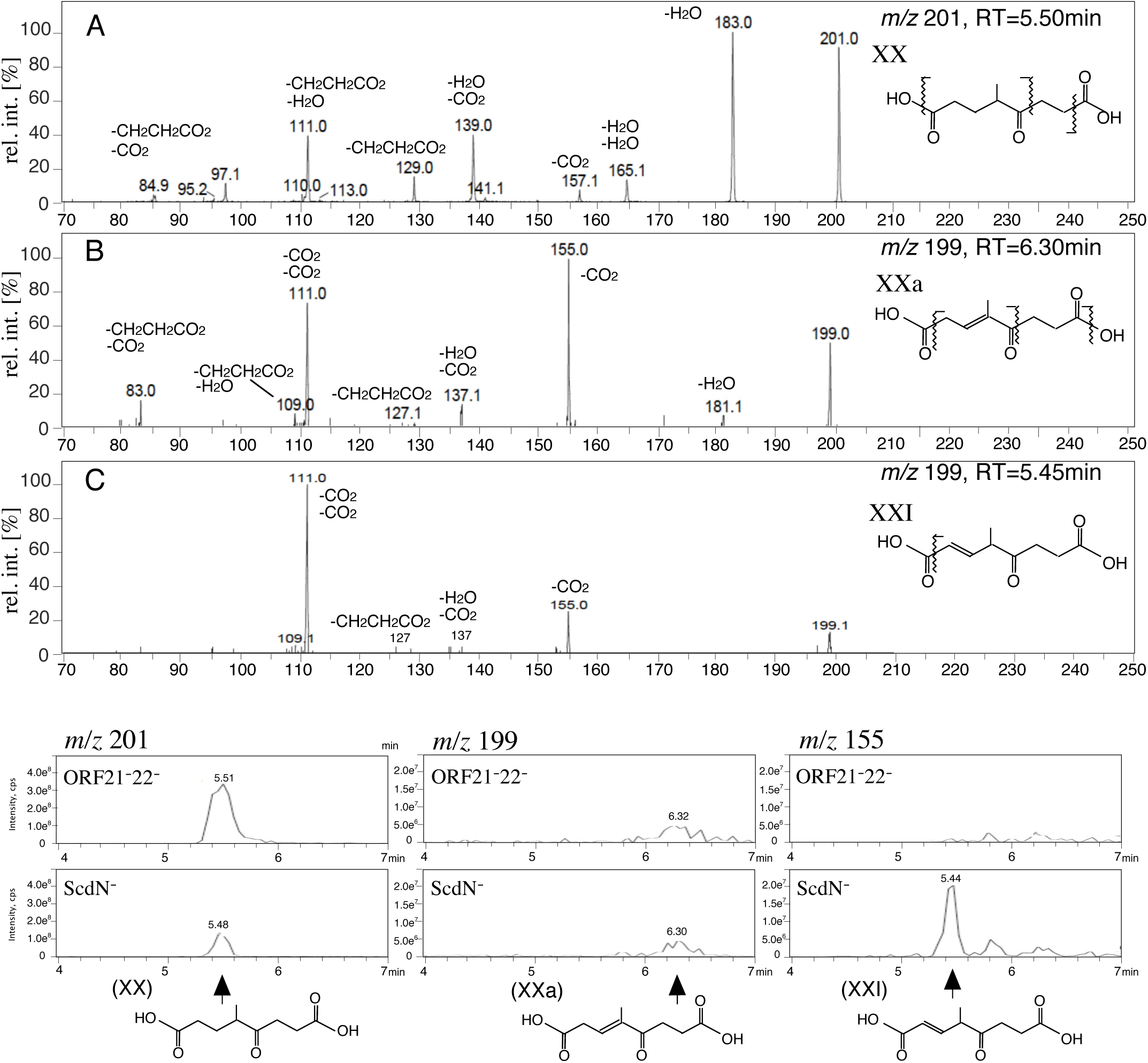
The mass spectra of compounds **XX** (*m/z* 201, A), **XXa** (*m/z* 199, B), and **XXI** (*m/z* 199, C) analyzed by LC/MS/MS. The mass chromatogram (*m/z* 201; **XX**, *m/z* 199; **XXa**, and *m/z* 155; **XXI**) are the same data as those in Fig. S1A to C in supplementary materials. Compounds are; 4-methyl-5-oxo-octane-1,8-dioic acid (**XX**), methyl-5-oxo-oct-3-ene-1,8-dioic acid (**XXa**), and 4-methyl-5-oxo-oct-2-ene-1,8-dioic acid (**XXI**). The vertical axis indicates relative intensity (%) and the horizontal axis indicates mass (*m/z*) in mass spectra. In mass chromatogram, the vertical axis indicates intensity (count/sec) and the horizontal axis indicates RT (min).

#### Complementation experiments using ORF21 and 22

Since ORF21^-^ and ORF22^-^ mutants accumulated significant amounts of **XX** and the homology search indicated that ORF21- and 22- encoded enzymes are similar to PimC and PimD, respectively. Therefore, ORF21 and ORF22 were expected to encode the large and the small subunit of the acyl-CoA dehydrogenase for **XX**-CoA ester. To confirm this, complementation experiments were carried out using ORF21^-^22^-^ mutants. An ORF21^-^22^-^ mutant carrying pMFY42 (negative control), ORF21^-^22^-^ mutant carrying pMFYORF21, ORF21^-^22^-^ mutant carrying pMFYORF22, and ORF21^-^22^-^ mutant carrying a pMFY42-based plasmid encoding ORF2122 (pMFYORF2122) were constructed and incubated with cholic acid (Fig. 4A-D).

**Fig 4.**
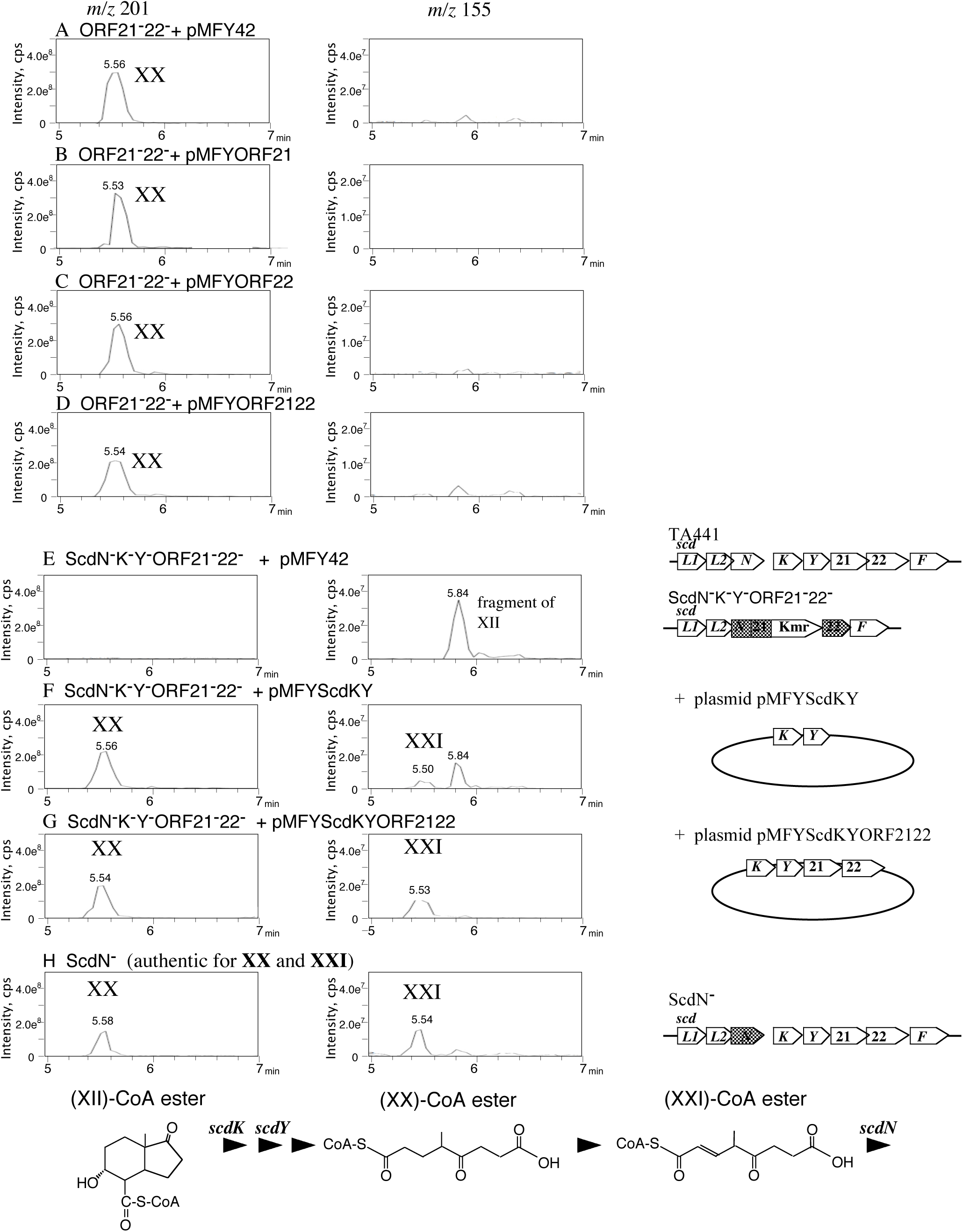
Complementation experiments with ORF21^-^22^-^ mutants. The mass chromatogram of each mutant culture incubated with 0.1% cholic acid for 7 d are shown. Mutants are; ORF21^-^22^-^ mutant carrying pMFY42 (ORF21^-^22^-^ with pMFY42) (A), carrying pMFYORF21 (B), carrying pMFYORF22 (C), carrying pMFY42 encoding ORF21 and ORF22 (ORF21^-^ with pMFYORF2122) (D), ScdN^-^,K^-^,Y^-^ORF21^-^22^-^ mutant carrying pMFY42 (ScdN^-^K^-^Y^-^ORF21^-^22^-^ with pMFY42) (E), carrying pMFY42 encoding *scdKY* (ScdN^-^K^-^Y^-^ORF21^-^22^-^ with pMFY*scdK,Y*) (F), carrying pMFY42 encoding *scdKY*ORF21,22 (ScdN^-^K^-^Y^-^ORF21^-^22^-^ with pMFY*scdKY*ORF21,22) (G), and ScdN^-^ mutant (as an authentic for **XX** and **XXI**) (H). Panels of *m/z* 211 and *m/z* 155 indicate **XX** and **XXI**, respectively. A peak with an *m/z* of 155 at RT = 0.58 min is the fragment of **XII**. Compounds are; 4-methyl-5-oxo-octane-1,8-dioic acid (**XX**), 4-methyl-5-oxo-oct-2-ene-1,8-dioic acid (**XXI**), and 9α-hydroxy-17-oxo-1,2,3,4,5,6,10,19-octanorandrostan-7-oic acid (**XII**). In mass chromatogram, the vertical axis indicates intensity (count/sec) and the horizontal axis indicates RT (min).

LC/MS/MS analysis demonstrated the accumulation of a smaller amount of **XX** in the culture of ORF21^-^22^-^ mutant carrying a pMFYORF2122 in comparison with the other three mutants, indicating that both ORF21 and ORF22 are indispensable for conversion of **XX**-CoA ester. However, the decrease in amount was small and the product, **XXI**, was scarcely detected (Fig. 4D *m/z* 155). The product is often detected in trace quantities in the complemented mutants, as all the steroidal degradation enzymes are expressed in the mutants [39–42].

To clearly detect **XXI** produced by ORF21- and 22-encoded enzyme, a new mutant in which *scdNKY*ORF21ORF22 was disrupted (ScdN^-^K^-^Y^-^ORF21^-^22^-^ mutant) carrying pMFY42, ScdN^-^K^-^Y^-^ORF21^-^22^-^ mutant carrying a pMFY42-based plasmid encoding *scdKY* (pMFY*scdKY*), and ScdN^-^K^-^Y^-^ORF21^-^22^-^ mutant carrying a pMFY42-based plasmid encoding *scdKY*ORF2122 (pMFY*scdKY*ORF2122) were constructed (Fig. 4E-G). Each mutant was incubated with cholic acid and the culture was analyzed by LC/MS/MS. A peak with an *m/z* of 155 at RT = 5.84 min is the fragment of 9α-hydroxy-17-oxo-1,2,3,4,5,6,10,19-octanorandrostan-7-oic acid (**XII,** Molecular weight 212, Fig. 4), which is an intermediate compound produced before **XV**. The mass chromatogram of the culture of a ScdN^-^ mutant is presented as authentic for **XXI** (Fig. 4H).

**XX** was not detected in the culture of ScdN^-^K^-^Y^-^ORF21^-^22^-^ mutant carrying pMFY42; however, a considerable amount of **XX** was accumulated in the culture of ScdN^-^K^-^Y^-^ORF21^-^22^-^ mutant carrying pMFY*scdKY* (Fig. 4E and 4F *m/z* 201). Considerable amount of **XXI** was detected only in the culture of the ScdN^-^K^-^Y^-^ORF21^-^22^-^ mutant carrying pMFY*scdKY*ORF2122, indicating the conversion of **XX**-CoA ester to **XXI**-CoA ester by ORF21- and 22-encoded enzymes. Small amount of **XXI** detected in the culture of the ScdN^-^K^-^Y^-^ORF21^-^22^-^ mutant carrying pMFY*scdKY* implies the conversion by other CoA-dehydrogenase as **XX** is a linear fatty acid (cf. ScdF and ScdJ described later); however, the difference in amount is considered to be the result of the conversion by the ORF21- and 22-encoded enzymes. Therefore, ORF21 and 22 were named *scdM1* and *scdM2,* respectively. ScdM1 and ScdM2 show homology to FadE31 (approximately 50 % amino acid identity) and FadE32 (approximately 30 % amino acid identity), respectively in *Mycobacterium tuberculosis* H37Rv. FadE31 and FadE32 in *M. tuberculosis* H37Rv were reported to form a dehydrogenase for **XX**-CoA ester because the amount of **XX**-CoA ester decreased upon treatment with purified FadE31 and FadE32 [45]. The function was confirmed only by the decrease in the amount of **XX**-CoA ester added as the substrate and the gene disruption was not performed in H37Rv.

### Structural elucidation of the CoA-esters involved in the degradation of steroidal C-ring (II); analysis of compounds XV, XVI, and XVII with LC/MS/MS

In the previous studies on C,D-ring cleavage, 9-oxo-1,2,3,4,5,6,10,19-octanor-13,17-secoandrost-8(14)-ene-7,17-dioic acid (**XV**)-CoA ester was thought to be the substrate of ScdL1L2 because the amount of **XV** and its derivatives (**XVa** and **XVb** in Fig. 1) was the largest among the compounds isolated from the culture of a ScdL1^-^L2^-^ mutant (Fig. 1, **XV, XVa, XVb, XVIa**, and **XVIIb**, all these five compounds were isolated and identified using NMR, HRMS, and other analysis when necessary [40]). Based on these compounds, **XV**-CoA ester, **XVI**-CoA ester, and **XVII**-CoA ester were proposed as intermediate compounds with only the C-ring remaining in bacterial steroid degradation (Fig. 1). ScdL1L2 shows the highest homology to acetate/3-ketoacid CoA transferases, which belong to the SugarP_isomerase superfamily (BLAST search). As the appropriate substrate of ScdM1M2 is **XX**-CoA, the intermediate compound produced just before **XX**-CoA was expected to be 6-methyl-3,7-dioxo-decane-1,10-dioic acid (**XIX**)-CoA ester. Therefore, the C-ring would be cleaved at C8-14. If **XV**-CoA ester is the substrate of CoA transferase ScdL1L2, **XV**-CoA ester is more likely to be cleaved at C8-9 than at C8-14. For cleavage at C8-14, the hydrated compound 14-hydroxy-9-oxo-1,2,3,4,5,6,10,19-octanor-13,17-secoandrostan-7,17-dioic acid **(XVIII)**-CoA ester is the most likely potential intermediate compound. **XVIII** was not isolated from the culture of the ScdL1^-^L2^-^ mutant in the previous study; however, **XV** and **XVII** might be the derivatives because the isolation was performed under acidic conditions. **XVIII** might be detected using LC/MS/MS as **XXI** was successfully identified; therefore, we analyzed the culture of the ScdL1^-^L2^-^ mutant using LC/MS/MS and obtained the MS spectrum of **XV**, **XVa**, **XVI**, **XVIa**, **XVII**, and **XVIIa** to obtain certain hints for the identification of **XVIII.**

**XVa** was detected as a large peak with an *m/z* of 181 at RT = 6.90 min (Fig. S1D). The mass spectrum showed fragments with *m/z* values of 137 and 109 (Fig. 5A). **XV** was detected as a faint peak with an *m/z* of 225 at RT = 6.30 min (Fig. S1E) with fragments with *m/z* values of 181, 163, 137, and 109 (Fig. 5B). **XVIa** was detected as a distinct fragment with an *m/z* of 197 at RT = 3.10 min (Fig. S1F) and the mass spectrum showed fragments with *m/z* values of 179, 153, 135, and 107 (197 and 135 were major) (Fig. 5C). In contrast to **XV**, **XVI** was detected as a distinctive fragment with an *m/z* of 241 at RT = 2.65 min (Fig. S1G) and the mass spectrum showed fragments with *m/z* values of 197, 179, 153, 135, and 107 (241, 197, and 135 were major) (Fig. 5D). **XVIIa**, the decarboxylated derivative of **XVII**, was detected as a peak with an *m/z* of 183 at RT = 7.00 min (Fig. S1H). The detected **XVIIa** was as abundant as **XVa**. In the previous study, **XVIIa** was not isolated because of the low UV absorption and only a small amount of **XVIIb** was isolated [40]; therefore, we presumed that the amount of **XVII**-CoA ester in the culture of the ScdL1^-^L2^-^ mutant was low. However, LC/MS analysis indicated the possibility that **XVII**-CoA ester is either one of the major intermediate compounds or its derivative. The mass spectrum of **XVIIa** showed a major fragment with an *m/z* of 139 (Fig. 5E). A peak with an *m/z* of 227 was detected at RT = 6.00 min (Fig. S1I), whose mass spectrum showed fragments with *m/z* values of 209, 183, 139, and 111, indicating that this peak is **XVII**. The mass spectral analysis showed that compounds with the C-ring are detected with the fragments “*m/z* 139 and 111”, “*m/z* 137 and 109”, or “*m/z* 135 and 107” (Fig. 5F, *m/z* 111 is not visible, but it was detected with higher collision energy).

**Fig 5.**
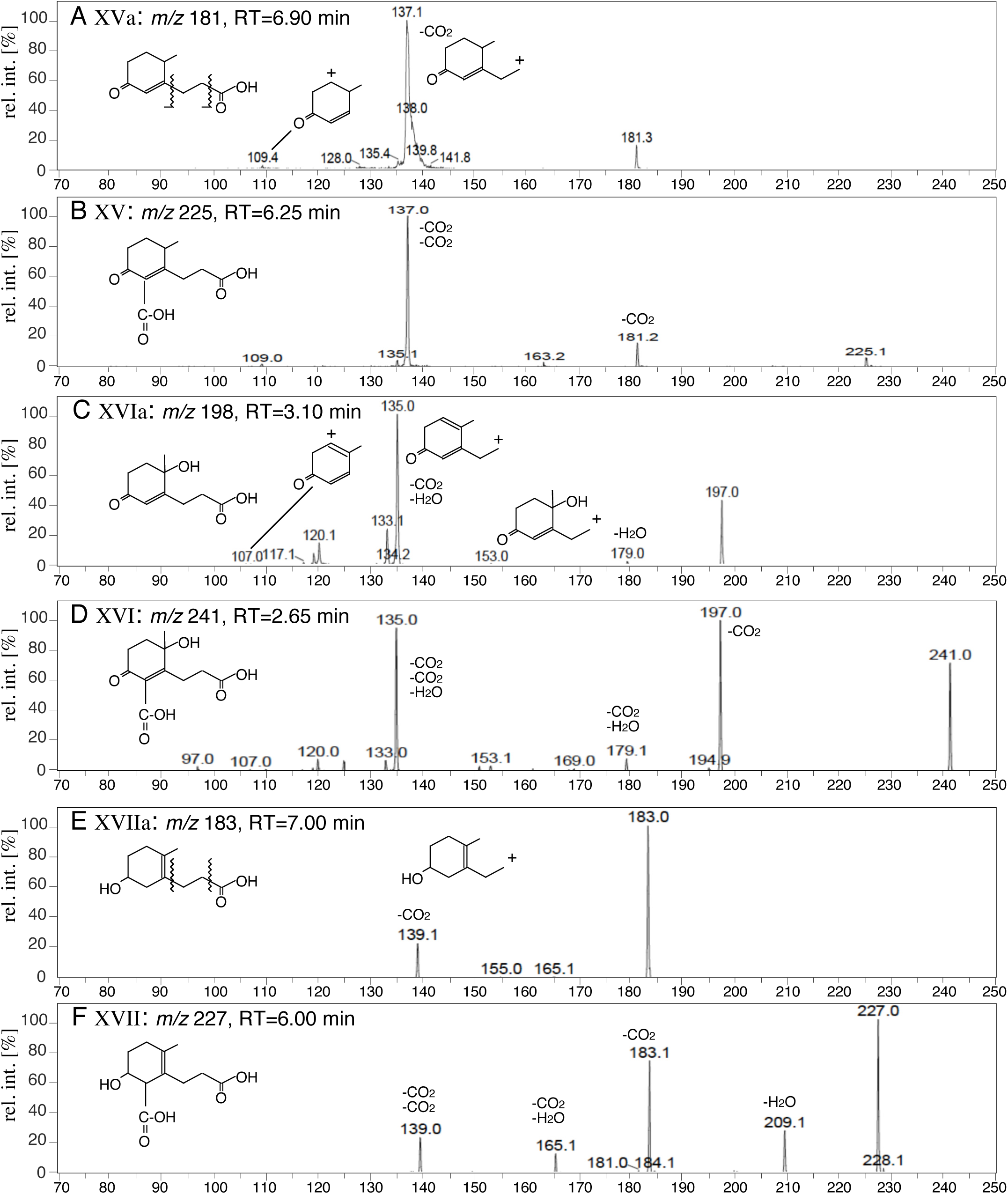
The mass spectra of the intermediate compounds in the culture of ScdL1^-^L2^-^ mutant incubated with cholic acid. Compounds are; 9-oxo-1,2,3,4,5,6,10,19-octanor-13,17-secoandrost-8(14)-en-17-oic acid (**XVa**) (A), 9-oxo-1,2,3,4,5,6,10,19-octanor-13,17-secoandrost-8(14)-ene-7,17-dioic acid (**XV**) (B), 13-hydroxy-9-oxo-1,2,3,4,5,6,10,19-octanor-13,17-secoandrost-8(14)-en-17-oic acid (**XVIa**) (C), 13-hydroxy-9-oxo-1,2,3,4,5,6,10,19-octanor-13,17-secoandrost-8(14)-ene-7,17-dioic acid (**XVI**) (D), 9-hydroxy-1,2,3,4,5,6,10,19-octanor-13,17-secoandrost-13-en-17-oic acid (**XVIIa**) (E), and 9-hydroxy-1,2,3,4,5,6,10,19-octanor-13,17-secoandrost-13-ene-7,17-dioic acid (**XVII**) (F). The mass chromatogram of these compounds are shown in Fig. S1D to S1I. The vertical axis indicates relative intensity (%) and the horizontal axis indicates mass (*m/z*).

### Compound XVIII

Molecular weight of **XVIII** is 244. In the mass chromatogram of *m/z* 243, a small peak at RT = 1.80 min was detected that was larger in amount in the culture of the ScdL1^-^L2^-^ mutant than in cultures of other mutants (Fig. S1J). The mass spectrum of this peak showed fragments with *m/z* values of 225, 119, 181, 171, 155,153, 137, 113, and 109, which suggested the presence of a C-ring with a ketone moiety and a double bond as a decomposed fragment (Fig. 6). The fragment with *m/z* values of 153 implied the presence of -CH_2_CH_2_CO_2_H and -OH and the fragment with *m/z* values of 155 implied the presence of two carboxylic acid groups. The mass spectral data was consistent with the structure of **XVIII**. In addition, a smaller peak with *m/z* of 243 at RT = 2.80 min (Fig. 8 *m/z* 243 B and Fig. S1J ScdL1^-^L2^-^ and ScdL1^-^L2^-^ + pMFY42), which is often detected with a peak with *m/z* of 243 at RT = 1.80 min, had a similar mass spectrum to that of the peak with *m/z* of 243 at RT = 1.80 min. This might be an isomer of **XVIII;** however, the mass spectrum of this compound was unclear because of the small quantity available in the samples. In our previous study, **XV** and its derivatives were isolated as the major compounds accumulated in the culture of ScdL1^-^L2^-^ mutant [40], while **XVIII** was not identified. **XVIII** was thought to be dehydrated at C-14 during the isolation and also during sample preparation for UPLC/MS and LC/MS/MS analyses, since both are carried out under acidic conditions. In the report on C,D-ring degradation by *M. tuberculosis* H37Rv [45], Crowe et al. described that **XV**-CoA ester was isolated and was identified using NMR analysis as the substrate of C-ring cleavage. We wanted to compare their NMR data with our NMR data of **XV** in order to obtain some indications that would help identify the substrate of ScdL1L2, however the NMR data of **XV**-CoA was not reported. In fact, Crowe et al. compared the major signals of **XV**-CoA with those of **XVa** presented in our Japanese patent in 2002 for identification of **XV**-CoA ester [45]. In their next paper, they reported that IpdAB of *Rhodococcus Jostii* RHA1 (similar to IpdAB of M. *tuberculosis* H37Rv and corresponds to ScdL1L2 of TA441) has both hydration activity at C 8-14 and isomeration activity to produce **XIX**-CoA ester from **XV**-CoA ester under the presence of FadA6, a thiolase corresponds to ScdF in TA441 [53]. However, we were not able to find the evidence for the hydration activity with ScdL1L2. Therefore, we carried out some more experiments to examine the substrate of ScdL1L2.

**Fig 6.**
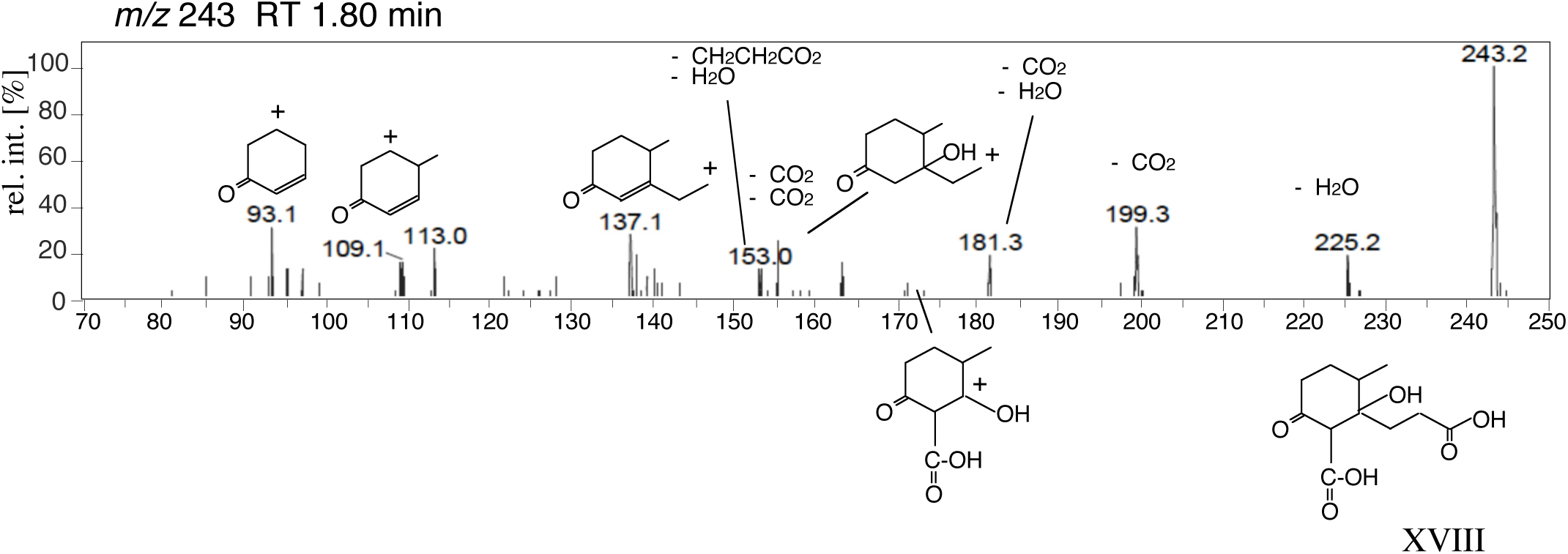
The mass spectrum of a peak with an *m/z* of 243 at RT = 1.80 min detected in the culture of ScdL1^-^L2^-^ mutant incubated with cholic acid. The mass chromatogram is shown in Fig. S1J. **VIII**: 14-hydroxy-9-oxo-1,2,3,4,5,6,10,19-octanor-13,17-secoandrostane-7,17-dioic acid. The vertical axis indicates relative intensity (%) and the horizontal axis indicates mass (*m/z*).

#### Complementation experiments using ScdL1L2 and the LC/MS/MS analysis

We carried out complementation experiments using ScdL1^-^L2^-^ mutant carrying a pMFY42-based and ScdL1^-^L2^-^ mutant carrying a pMFY42-based plasmid coding ScdL1L2 (pMFYScdL1L2) and analyzed the culture with LC/MS/MS to examine whether **XVIII** was converted in the mutant expressing ScdL1L2. In the culture of the ScdL1^-^L2^-^ mutant carrying pMFYScdL1L2, **XVIII** was slightly less abundant than ScdL1^-^L2^-^ mutant carrying a pMFY42, though the peaks were too small to confirm the conversion (Fig. 7A and B). This is probably due to the insufficient expression of ScdL1L2 from pMFYScdL1L2, as observed with ScdM1M2.

**Fig. 7.**
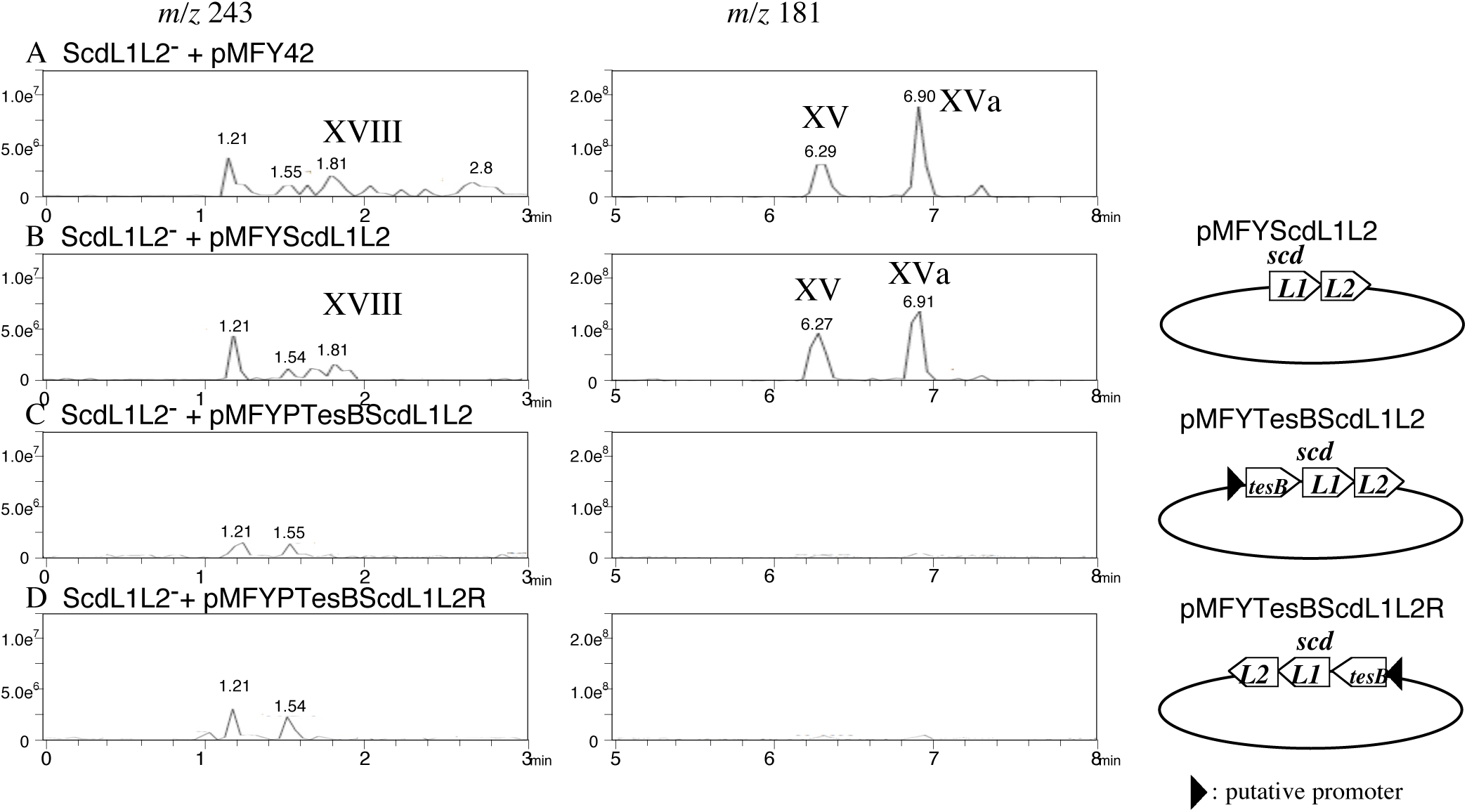
Complementation experiments with ScdL1^-^L2^-^ mutants. The mass chromatogram of each mutant culture incubated with 0.1% cholic acid for 7 d are shown. Mutants are; ScdL1^-^L2^-^ mutant carrying pMFY42 (ScdL1^-^L2^-^ with pMFY42) (A), carrying pMFYScdL1L2 (pMFY42-derivative carrying *scdL1L2*) (ScdL1^-^L2^-^ with pMFYscdL1L2) (B), carrying pMFYTesBScdL1L2 (pMFY42-derivative carrying the putative promoter region and *tesBscdL1L2* in the same direction of the genes on pMFY42) (ScdL1^-^L2^-^ with pMFYtesBscdL1L2) (C), and carrying pMFYTesBScdL1L2R (pMFY42-derivative carrying the putative promoter region and *tesBscdL1L2* in the opposite direction of the genes on pMFY42) (ScdL1^-^L2^-^ with pMFYtesBscdL1L2R) (D). Panels of *m/z* 243 indicate **XVIII** and panels of *m/z* 181 indicate **XV** and **XVa**. Compounds are; 14-hydroxy-9-oxo-1,2,3,4,5,6,10,19-octanor-13,17-secoandrostane-7,17-dioic acid (**VIII**), 9-oxo-1,2,3,4,5,6,10,19-octanor-13,17-secoandrost-8(14)-ene-7,17-dioic acid (**XV**), and 9-oxo-1,2,3,4,5,6,10,19-octanor-13,17-secoandrost-8(14)-en-17-oic acid (**XVa**). In mass chromatogram, the vertical axis indicates intensity (count/sec) and the horizontal axis indicates RT (min).

Therefore, we constructed two more mutants; ScdL1^-^L2^-^ mutant carrying a pMFY42 coding *tesBscdL1L2* (pMFYTesBScdL1L2) in the same direction as the plasmid genes (Fig. 7C) and in the opposite direction (pMFYTesBScdL1L2R) (Fig. 7D). These plasmids contain the putative promoter of steroid degradation genes *tesB* to *tesR* in the upstream DNA region of *tesB*, so the transcription level of *scdL1L2* is expected to be higher in these mutants than in ScdL1^-^L2^-^ mutant carrying pMFYScdL1L2. TesB is the *meta-*cleavage enzyme for cleavage of the aromatized A-ring; therefore, *tesB* on the plasmids does not affect C,D-ring degradation (cf. Fig. 1). These mutants were cultured in the same way as the ScdL1^-^L2 ^-^ mutant carrying pMFYScdL1L2 and the culture was analyzed using LC/MS/MS. **XVIII**, **XV**, and **XVa** were almost undetectable in the culture with these mutants. The results indicated that ScdL1L2 expressed from pMFYTesBScdL1L2 and pMFYTesBScdL1L2R in these mutants had enough activity as that in the wild type (Fig. 7C and 7D). However, other characteristic quantitative change of intermediate compounds to suggest the actual substrate of ScdL1L2 was not detected. ScdL1L2 showed highest homology to acetate/3-ketoacid CoA transferases, which belong to the SugarP_isomerase superfamily (NCBI BLAST search). Therefore, function of ScdL1L2 in steroid degradation is likely to be the isomerization of **XVIII**-CoA ester to **XIX**-CoA ester. The results of the two mutants, ScdL1^-^L2^-^ mutant carrying pMFYTesBScdL1L2 and ScdL1^-^L2^-^ mutant carrying pMFYTesBScdL1L2R were almost the same, indicating that the enzyme expression from plasmid promoter is quite low compared to the expression from the promoter of steroid degradation genes *tesB* to *tesR*, which resulted in the insufficient conversion in complemented mutants.

### Conversion of XVIII-CoA ester by crude extract containing ScdL1L2

**XX**-CoA ester is hypothesized to be produced from 6-methyl-3,7-dioxo-decane-1,10-dioic acid (**XIX**)-CoA ester by acyl-CoA transferase reaction (Fig. 1). However, **XIX** (molecular weight 244) was not identified in previous studies. Mass chromatogram of *m/z* 243 implied the possibility that a compound detected at RT = 1.55 min might be **XIX** (Fig. S1J), but the detected amount was too small for identification (cf. Fig. S1, mass spectrum of the peak with *m/z* 243 at RT = 1.55 min). Therefore, we attempted the conversion of **XVIII**-CoA ester in the culture of the ScdL1^-^L2^-^ mutant with *Escherichia coli* expressing ScdL1L2. The supernatant of the culture of ScdL1^-^L2^-^ mutant after cell disruption was used as the substrate solution as isolating the **XVIII**-CoA ester was difficult. *E. coli* carrying a pUC19-based plasmid expressing ScdL1L2 or pUC19 (negative control) was independently cultured and the cells were disrupted by sonication in the substrate solution on ice, after which the samples were incubated at 30°C. The LC/MS analysis showed a slight decrease in **XVIII** abundance in ScdL1L2-treated samples compared to the negative control (Fig. 8, *m/z* 243), however an increase in the amount of **XIX** and decrease of XV (Fig. 8, *m/z* 225 and *m/z* 181 5 min-8 min) was not detected. Instead, a clear peak with *m/z* of 181 at RT = 3.35 min was detected only in the samples treated with *E. coli* expressing ScdL1L2 (Fig. 8, *m/z* 181 1 min-4 min). We could not identify this peak successfully; however, the mass spectrum of this peak showed fragments with *m/z* values of 163 and 137 (Fig. 8D), which suggested the presence of a carboxyl group and a hydroxyl group. Fragments with *m/z* values of 181 and 137 are often detected in the mass spectrum of intermediate compounds with a C-ring, yet compounds with a molecular weight of 181 with the C-ring were detected at an RT of approximately 6−7 min (Fig. 5A, 7 *m/z* 181, and S1D). Therefore, the compound detected as a peak with *m/z* of 181 at RT = 3.35 min was not likely to have any rings. In addition to this compound, **XX** increased in amount in the samples treated with *E. coli* expressing ScdL1L2 (Fig. 8, *m/z* 201). Since **XIX** is a linear fatty acid and we used the disrupted cells of *E. coli* expressing ScdL1L2 for this conversion experiment, some CoA-transferases from TA441 cells (and possibly *E. coli* cells) were thought to have converted **XIX**-CoA-ester to **XX**-CoA-ester (A similar conversion was observed in the complementation experiment with ScdM1M2, Fig. 4F).

**Fig. 8.**
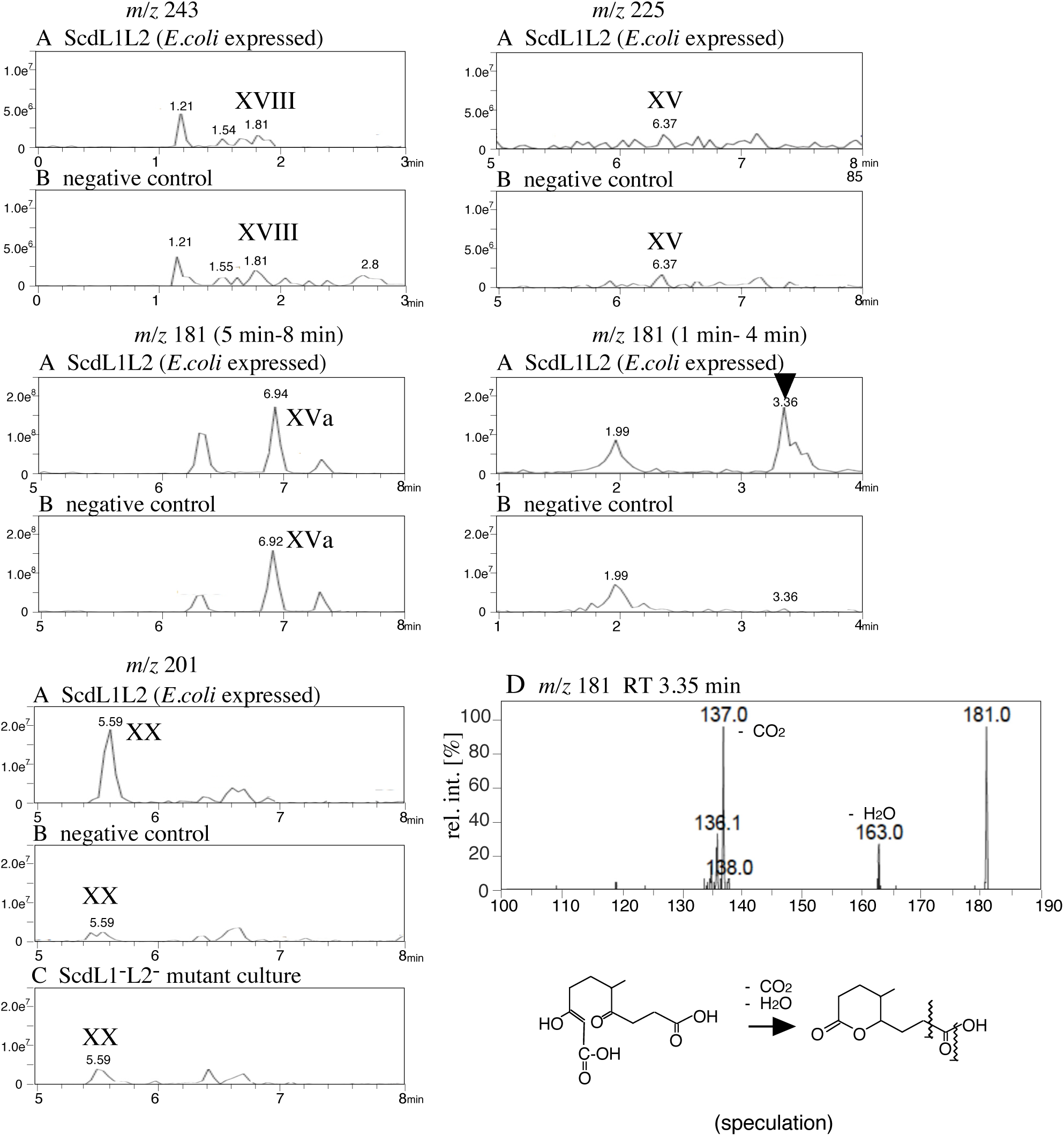
Conversion experiment using the enzyme ScdL1L2 expressed in *E.coli*. The mass chromatogram of the reaction solution containing **XIII**-CoA ester treated with ScdL1L2 (A) and without ScdL1L2 (B) are shown. Panel C shows the mass chromatogram of the ScdL1^-^L2^-^ mutant culture to confirm that **XX** detected in panel B (*m/z* 201) was contained in the reaction solution. The vertical axis indicates intensity (count/sec) and the horizontal axis indicates RT (min). Panel D shows the mass spectrum of the peak with an *m/z* of 181 at RT = 3.35 min. The vertical axis indicates intensity (count/sec) in mass chromatogram and relative intensity (%) in mass spectra, and the horizontal axis indicates mass (*m/z*). Compounds are; 14-hydroxy-9-oxo-1,2,3,4,5,6,10,19-octanor-13,17-secoandrostane-7,17-dioic acid (**VIII**) and 4-methyl-5-oxo-octane-1,8-dioic acid (**XX**).

### CoA-transferase, ScdF

Our previous study showed that the primary role of ScdF is that of CoA-transferase for conversion of **XI**-CoA ester to **XII**-CoA ester (Fig. 1) [42]. In *M. tuberculosis* H37Rv, FadA6, the enzyme that corresponds to ScdF, was reported as a CoA-transferase, which converts **XIX**-CoA-ester to **XX**-CoA-ester [45]. Moreover, IpdAB of *M*. *tuberculosis* H37Rv (corresponds to ScdL1L2 of TA441) shows hydration activity at C 8-14 and isomeration activity to produce **XIX**-CoA ester from **XV**-CoA ester only under the presence of FadA6, a thiolase corresponds to ScdF in TA441 [53]. Since the previous study suggested that a peak with *m/z* of 143 detected in the culture of ScdF^-^ mutant might be the fragment of the product of **XVIII**-CoA ester, we analyzed the mass chromatograms and the mass spectrum for peaks with *m/z* of 143 in the several mutant cultures with LC/MS/MS. The analysis showed a larger peak with an *m/z* of 143 at RT = 3.00 min in the cultures of the ScdF^-^ mutant, ScdM1^-^M2^-^ mutant, and ScdN^-^ mutant than in the cultures of the ScdY^-^ mutant and ScdL1^-^L2^-^ mutant (Fig. S1K). However, the mass spectrum of the peak did not provide any clues to speculate the structure of the compound. We then analyzed the cultures of ScdF^-^ mutant and ScdJ^-^ mutant incubated with testosterone to examine the involvement of ScdF and ScdJ in conversion of **XI**-CoA ester to **XII**-CoA ester (Fig. S2). Testosterone was used because the results would be complicated with cholic acid which has a hydroxyl group at C-7α; optical isomers □□□□ (with 7α-OH) and **X** (with 7β-OH) are produced in the degradation of cholic acid (Fig. 1). Large amount of **XI**, **XIa** (the lactone form **XI**), **X,** and **Xa** (the lactone form **X**) were detected in the ScdF^-^ mutant culture (**XI**-CoA ester is produced from **X**-CoA ester) (Fig. S2A and Fig. S2C), while negligible amount of **XI**, **XIa**, and **Xa** with a small amount of **X** was detected in the ScdJ^-^ mutant culture. All the results we obtained on ScdF and ScdJ supported that the primary role of ScdF and ScdJ is that of CoA-transferases for conversion of **XI**-CoA ester to **XII**-CoA ester and that of **XIX**-CoA ester to **XX**-CoA ester.

In conclusion, this study found the degradation pathway of the steroidal C-ring as shown in Fig. 9. **XVIII**-CoA is likely to be the suitable substrate of ScdL1L2 and converted to **XIX**-CoA, which is followed by the conversion of **XIX**-CoA to **XX**-CoA mainly by ScdJ. ScdF may be involved in the reaction; however, the primary role of ScdF is conversion of **XIX**-CoA ester to **XX**-CoA ester. Then, **XX**-CoA ester was dehydrogenated by the ScdM1M2-encoded CoA-dehydrogenase to **XXI**-CoA ester, and a water molecule was added by ScdN for further degradation by β-oxidation. ScdN is considered to catalyze the last reaction in C,D-ring degradation by the enzymes encoded in the steroid degradation gene cluster *tesB* to *tesR*. Compounds **XVI**, **XVII**, **XVIII**, and **XXI**, which were not directly detected and only the decarboxylated or dehydrated derivatives were identified in the previous studies, were confirmed with LC/MS/MS analysis in this study. The mechanism of production of **XVIII**-CoA in *C. testosteroni* still needs to be elucidated in order to gain a detailed understanding of the whole degradation pathway of steroidal C,D-ring.

**Fig. 9.**
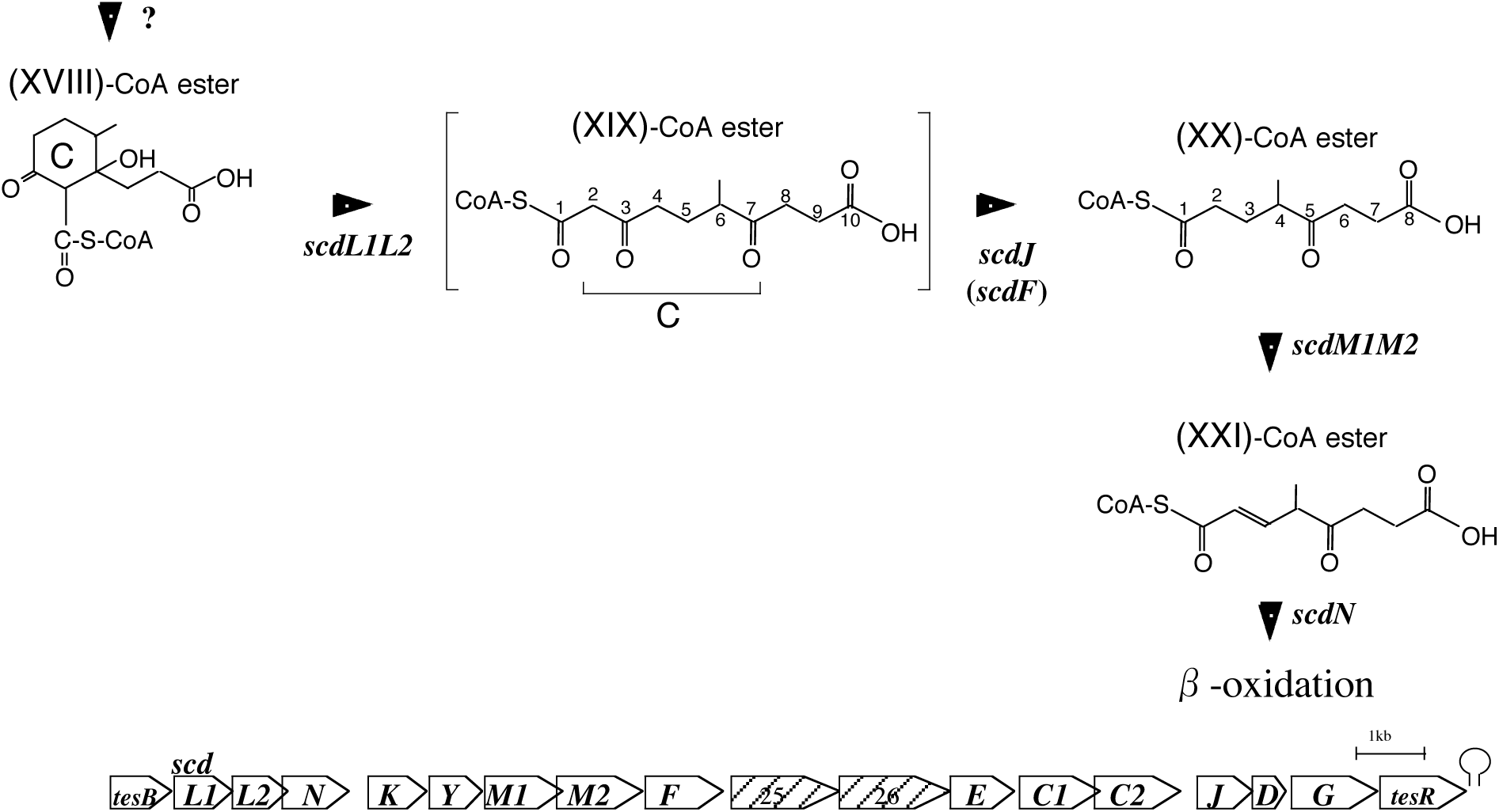
Intermediates and genes involved in the cleavage and degradation of the steroidal C-ring revealed in this study. Compounds are; 14-hydroxy-9-oxo-1,2,3,4,5,6,10,19-octanor-13,17-secoandrostane-7,17-dioic acid (**VIII**), 6-methyl-3,7-dioxo-decane-1,10-dioic acid (**XIX**), 4-methyl-5-oxo-octane-1,8-dioic acid (**XX**), and 3-hydroxy-4-methyl-5-oxo-oct-2-ene-1,8-dioic acid (**XXI**). Steroid degradation gene cluster *tesB* to *tesR* with genes revealed in this study (*scdM1M2* and *scdJ*) is shown below.

## MATERIALS AND METHODS

### Abbreviations

HPLC, high-performance liquid chromatography; UPLC/MS, ultra high-performance liquid chromatography-mass spectrometry; RT, retention time; MS, mass spectrometry; HRMS; high-resolution mass spectrometry, MW, molecular weight; CoA, coenzyme A.

### Culture conditions

Mutant strains of *C. testosteroni* TA441 were grown at 30°C in a mixture of equal volumes of Luria-Bertani (LB) medium and C medium (a mineral medium for TA441) [28, 54] with suitable carbon sources. This mixed media is used because the mutants accumulate more amount of intermediate compounds than with C medium or LB medium (unpublished data). Cholic acid and other steroids were added as filter-sterilized DMSO solutions with a final concentration of 0.1% (w/v).

### Construction of gene-disrupted mutants, plasmids, and mutants for complementation experiments

For construction of the ORF21^-^ mutant, ORF22^-^ mutant, ORF21^-^22^-^ mutant, and ORF33^-^ mutant (Table 1), pUC19 [55]-based plasmid carrying DNA region of TA441 with insertion of a kanamycin-resistance gene (Km^r^) without a terminator in the *Acc*I site in ORF21 (pUCORF21-Km^r^), in the *Hinc*II site in ORF22 (pUCORF22-Km^r^), between the *Apa*I site in ORF21 and *Apa*I site in ORF22 (pUCORF2122-Km^r^), and in the *Cla*I site in ORF33 (pUCORF33-Km^r^) were used (Table 2). For ScdN^-^K^-^Y^-^ORF21^-^22^-^ mutant, DNA fragment containing disrupted ORF2122 (PCR amplified using pUCORF2122-Km^r^ as the template DNA) was transferred into the pUCscdL1L2d (pUC19-based plasmid carrying 3.2 kb *Pst*I-*Pvu*II fragment containing *scdL1L2* and partial *scdN*) just after the 3.2 kb fragment to construct pUCscdN-ORF22-Km^r^ (Table 2). These plasmids were used for inactivation of the objective genes in TA441 by homologous recombination. The plasmids were introduced into TA441 via electroporation and Km^r^ colonies were selected [54]. Insertion of the Km^r^ was confirmed by southern hybridization and/or PCR-amplification. DNA fragments containing ORF21, ORF22, ORF2122, ORF33, *scdKY*, and *scdKY*ORF2122 were obtained by PCR amplification and introduced into a broad-host-range plasmid pMFY42 [56], which can be maintained in *Pseudomonas spp.* and several related species conferring tetracycline resistance to construct pMFYORF21, pMFYORF22, pMFYORF2122, pMFYORF33, pMFY*scdKY*, and pMFY*scdKY*ORF2122, respectively. Retention of the plasmids by the gene-disrupted mutants and the transformants was confirmed by Southern hybridization and/or PCR-amplification. For PCR amplification, DNA polymerase KOD -plus- ver. 2 (TOYOBO, Japan) was used. The primers are listed in Table 3.

**Table 1.** strains

| strains | Characteristics | Source or reference |
| --- | --- | --- |
| TA441 | Wild-type | [54] |
| ScdN<sup>-</sup> | *ScdN*: :Km<sup>r</sup> mutant (*Bam*HI site) of TA441 | [41] |
| ScdF<sup>-</sup> | *ScdF*: :Km<sup>r</sup> mutant (*Hinc*II site) of TA441 | [42] |
| ScdL1<sup>-</sup>L2<sup>-</sup> | *ScdL1L2*: :Km<sup>r</sup> mutant (*Apa*I site to *Eco*T22I site) of TA441 | [40] |
| ScdC1<sup>-</sup>C2<sup>-</sup> | *ScdC1C2*: :Km<sup>r</sup> mutant (*Eco*T22I site to *Nru*I site) of TA441 | [37] |
| ORF21<sup>-</sup> | ORF21: :Km<sup>r</sup> mutant (*Acc*I site) of TA441 | this work |
| ORF22<sup>-</sup> | ORF22: :Km<sup>r</sup> mutant (*Hinc*II site) of TA441 | this work |
| ORF21<sup>-</sup>22<sup>-</sup> | ORF2122: :Km<sup>r</sup> mutant (*Apa*I site) of TA441 | this work |
| ORF33<sup>-</sup> | ORF33: :Km<sup>r</sup> mutant (*Cla*I site) of TA441 | this work |
| ScdN<sup>-</sup>K<sup>-</sup>Y<sup>-</sup>ORF21<sup>-</sup>22<sup>-</sup> | *ScdN* to ORF22: : Km<sup>r</sup> mutant of TA441 | this work |

Km<sup>r</sup>: Km-resistance, [54] Arai et al. 1998. Microbiology **144**: 2895-2903 [41] Horinouchi et al. 2001. Microbiology **147**:3367-3375 [42] Horinouchi et al. 2019. Appl Environ Microbiol **85**:e001204-19 [40] Horinouchi et al. 2019. J Steroid Biochem Mol Biol, **185**: 268-276 [37] Horinouchi et al. 2014. J Bacteriol **196**:3598-3608

**Table 2.** plasmids

| plasmids | Characteristics | Source or reference |
| --- | --- | --- |
| pUC19 | Ap<sup>r</sup>, *lacZ* | [55] |
| pMFY42 | Tc<sup>r</sup>, Km<sup>r</sup>, RSF1010-based broad host range plasmid | [56] |
| pCos15-21 | Ap<sup>r</sup>, pSuperCosI* with *Bam*HI insert of TA441 (containing *steA* to *tesR*) | [31] |
| pUCORF21d | pUC19 derivative carrying 2.3 kb *Hinc*II-fragment from pCos15-21 | this work |
| pUCORF21-Km<sup>r</sup> | pUCORF21d derivative with Km<sup>r</sup> in *Acc*I site of ORF21 | this work |
| pUCORF22d | pUC19 derivative carrying 3.8 kb *Eco*RI-*Hind*III fragment from pCos15-21 | this work |
| pUCORF22-Km<sup>r</sup> | pUCORF22d derivative with Km<sup>r</sup> in *Hinc*II site of ORF22 | this work |
| pUCORF2122d | pUC19 derivative carrying 4.5 kb *Kpn*I-*Hind*III fragment from pCos15-21 | this work |
| pUCORF2122-Km<sup>r</sup> | pUCORF2122d derivative with Km<sup>r</sup> in the region between *Apa*I site of ORF21 and *Apa*I site of ORF22 | this work |
| pUCORF33d | pUC19 derivative carrying 3.0 kb *Eco*RI-*Bgl*II fragment from pCos15-21 | this work |
| pUCORF33-Km<sup>r</sup> | pUCORF22d derivative with Km<sup>r</sup> in *Cla*I site of ORF33 | this work |
| pUCscdL1L2d | pUC19 derivative carrying 3.2 kb *Pst*I-*Pvu*II fragment of pCos15-21 (containing *scdL1L2*) in *Hinc*II site | this work |
| pUC scdN-ORF22-Km<sup>r</sup> | pUCscdL1L2d derivative carrying PCR amplified DNA fragment of ORF21<sup>-</sup>22<sup>-</sup> mutant genome (containing ORF2122: :Km<sup>r</sup> and *scdF*) in *Sma*I site in the downstream region of *scdL2* | this work |
| pMFYORF21 | pMFY42 derivative carrying ORF21 | this work |
| pMFYORF22 | pMFY42 derivative carrying ORF22 | this work |
| pMFYORF2122 | pMFY42 derivative carrying ORF2122 | this work |
| pMFYscdYKORF2122 | pMFY42 derivative carrying *scdYK* and ORF2122 | this work |
| pMFYscdYK | pMFY42 derivative carrying *scdYK* | this work |
| pMFYTesBScdL1L2 | pMFY42 derivative carrying the promoter region, *tesB*, and *scdL1L2* | this work |
| pMFYTesBScdL1L2f | pMFY42 derivative carrying the promoter region, *tesB*, and *scdL1L2* in the opposite direction of the pMFY42 genes | this work |

Km<sup>r</sup>: Km-resistance pSuperCosI\* (Stratagene, CA) [55] Vieira, J., and Messing, J. 1987. Methods Enzymol. **153**: 3-11. [56] Nagata Y. et al. 1993. J Bacteriol **175**:6403-6410 [31] Horinouchi M, et al. 2004. Biochem Biophys Res Commun **324**:597-604.

**Table 3.**
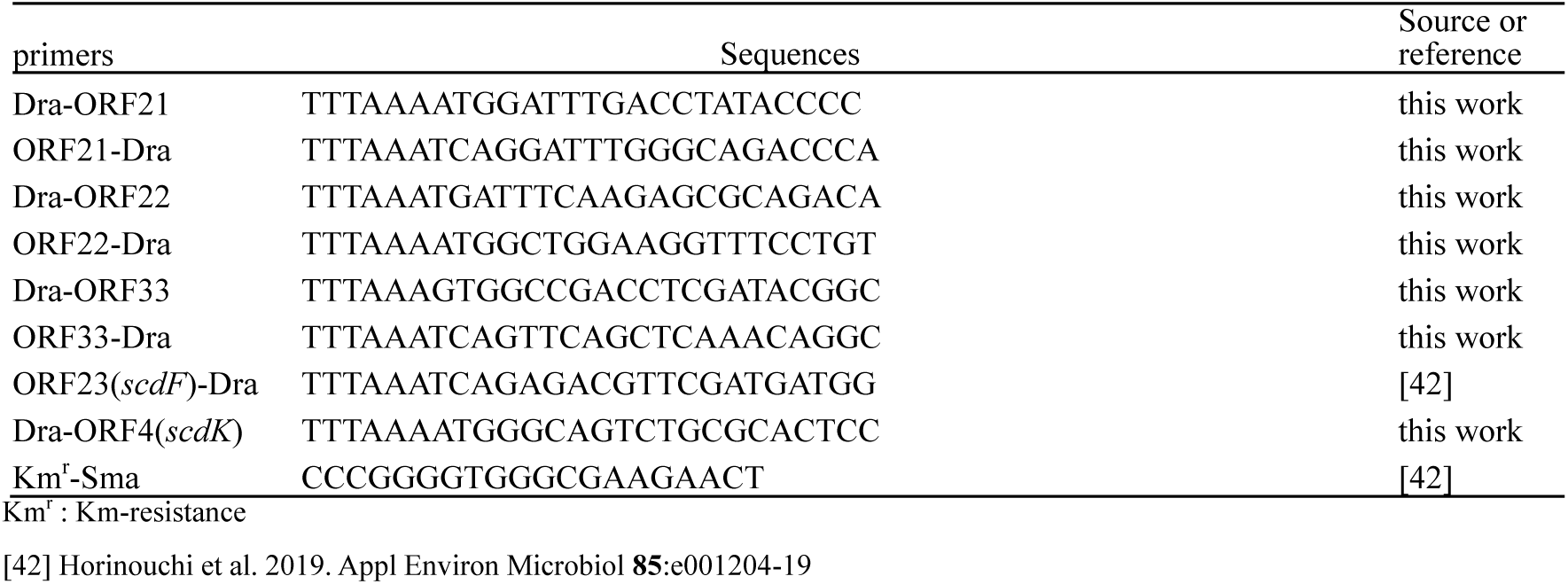
primers

### Conversion of XVIII-CoA ester by ScdL1L2

ScdL1^-^L2^-^ mutant was cultured in LB + C-medium with 0.1 % w/v cholic acid for 7 d. The cells were disrupted by freezing and thawing three times, followed by sonication (at 45 kHz, three bursts of 20 s each, with 1 min intervals in 50 mL tube) on ice. The culture was centrifuged in 1.5 mL tubes (15,000 rpm, 10 min, 4°C) and the supernatant was used as the substrate solution because isolation of the **XVIII**-CoA ester was difficult. *E. coli* carrying pUC19 derivative plasmid expressing ScdL1L2 and pUC19 (negative control) was independently cultured and the cells were collected by centrifugation in 1.5 mL tubes. One milliliter of the substrate solution was added to each microcentrifuge tube and the *E.coli* cells were disrupted by sonication (at 20 kHz, three bursts of 20 s each, with 1 min intervals) on ice. The whole procedure was carried out on ice or at 4°C, and then the samples were immediately incubated at 30°C for 1 h.

### Ultra high-performance liquid chromatography (UPLC)/MS

The 1ml-culture was extracted twice with a double volume of ethyl acetate under acidic conditions (pH 2 with HCl). The ethyl acetate layer was dried and dissolved in 550µl-methanol and the 1.5 µl of the methanol solution was injected to UPLC/MS. UPLC/MS was carried out using an Applied Biosystems Q Trap liquid chromatography tandem MS system with a reverse phase column (XTerra MSC_18_, 2.1 × 150 mm, Waters) at a flow rate of 0.4 ml/min at 40°C. Elution was carried out using a linear gradient from 20% solution A (CH_3_CN) and 80% solution B (H_2_O:HCOOH = 100:0.05) to 100% solution A over 4.5 min, which was maintained for 2 min. Electrospray ionization (negative) was used for the detection. The conditions for MS were an ion spray voltage of 5.0 kV, a curtain gas pressure of 15 psi, a nebulizer gas pressure of 40 psi, an auxiliary gas pressure of 60 psi, and an ion source temperature of 400°C.

### Reverse-phase liquid chromatography with tandem mass spectrometry (LC/MS/MS)

For LC/MS/MS analysis, 2 µl of the samples prepared the same way as those for HPLC/MS analysis was injected into the system. Agilent 1100 HPLC (Agilent, CA) with a mass spectrometer (Applied Biosystems 4000 Q-TRAP, MS) was used with L-column2 ODS (1.5 × 150 mm) Type L2-C 18.5µm, 12mm (GL Science, Tokyo, Japan) and elution was carried out using 90% solution C (H_2_O:HCOOH = 100:0.1) and 10% solution A for 1min, followed by a linear gradient from 90% solution C and 10% solution A to 20% solution C and 80% solution A over 7 min, which was maintained for 2 min. The flow rate was 0.2 ml/min. The desolvation temperature for the mass spectrometer was fixed at 450 °C. The collision energy was 20 V.

## ACKNOWLEDGMENTS

MH appreciates Dr. Reizo Kato (head of Condensed Molecular Materials Laboratory, RIKEN) and Dr. Yousoo Kim (head of Surface and Interface Science Laboratory, RIKEN) for thoughtful supports and advices. The authors thank Dr. Takemichi Nakamura (Molecular Structure Characterization Unit, RIKEN CSRS, WAKO) for his assistance in mass spectrometry, and Dr. Toshihiko Nogawa and Dr. Hiroyuki Osada (RIKEN CSRS) for use of UPLC/MS

